# Finger Type Classification for Fingerprint Image Error Correction in Large Scale Biometric Databases

**DOI:** 10.64898/2025.12.14.694185

**Authors:** Tahsin Islam. Sakif, Nasser Nasrabadi, Jeremy Dawson

## Abstract

Large-scale biometric systems, essential for national security and border management, increasingly rely on multimodal databases containing millions of identities. However, operational pressures and insufficient training lead to frequent image classification and labeling errors by human operators. These critical data integrity issues include the mislabeling of rolled vs. flat fingerprints, out-of-sequence captures, and the insertion of incorrect modalities. Such errors render enrollment records unreliable, compromising subsequent identity verification processes. Since manually sorting vast image archives is unfeasible, our study proposes an automated solution. The primary objective was to deploy a Siamese Network to classify fingerprints by their precise finger type and collection methodology (flat or rolled impressions). A secondary, but central, goal was to investigate the influence of varying embedding dimensions (64, 128, 256, 512) and similarity thresholds (0.5, 0.2, 0.1) on the network’s performance metrics. Our most significant finding demonstrates a clear trade-off: a lower similarity threshold drastically increases conditional accuracy and precision (e.g., up to 98%) but simultaneously increases the proportion of images categorized as "uncertain" (up to 24%). In a practical, large-scale application, this necessitates balancing superior classification accuracy against a higher volume of images requiring costly manual inspection. This work provides a proof-of-concept tool capable of efficiently quantifying the percentage of images requiring human review across various modalities (fingerprints, face, iris). The eventual goal is a lightweight, efficient tool to establish standard preprocessing procedures for any large biometric dataset, dramatically reducing the time and cost associated with data integrity maintenance.

## Introduction

The proliferation of large-scale biometric systems has established them as foundational components of modern civil identification, border control, and national security infrastructure. Multimodal databases, such as those maintained by the U.S. Department of Homeland Security (DHS) or the national Aadhaar system in India, house hundreds of millions of identity records, including face, iris, and fingerprint modalities, with volumes expanding rapidly [1-2]. This immense reliance on biometric data quality, however, is compromised by inherent challenges in data acquisition [3].

The central issue is the pervasive problem of data integrity failure stemming from human operational error during enrollment. High throughput demands, coupled with insufficient operator training, frequently result in critical image classification and labeling inaccuracies within these centralized archives. Concrete examples of these errors, documented in law enforcement and government systems (e.g., EBTS records), include: the incorrect sequencing of finger images, intra-modality misclassification (such as labeling a rolled fingerprint as a flat impression), and inter-modality errors (inserting a face or iris image into a fingerprint field) [4-6]. The presence of inaccurate data has been shown to degrade the reliability of automated systems, leading to high false rejection rates and the risk of wrongful identification or denial of service [4]. Consequently, ensuring the quality and correct classification of raw data is paramount for maintaining the efficacy of the entire biometric ecosystem.

Currently, the process of rectifying these enrollment errors commonly referred to as de-duplication or sequence checking remains a largely manual, tedious, and unscalable task [7-8]. A manual quality assurance process is simply unsustainable for databases containing millions of records. This research addresses this critical gap by developing an automated, deep learning solution for identifying misclassified images, thereby reducing the immense time and cost associated with human inspection [9-17].

The human ability to differentiate and classify fingerprints based on their unique ridge patterns provides the conceptual basis for our automated approach. Expert latent print examiners, through sufficient training, are demonstrably capable of sorting prints based on specific finger types [17]. Similarly, Convolutional Neural Networks (CNNs) are highly suitable for this task because fingerprints possess a specific, repetitive composition of minutiae and ridges [9-10]. Deep learning architectures can effectively learn the statistical characteristics of these patterns, offering a robust alternative to conventional minutiae matching techniques that are sensitive to noise and computational expense [9].

The goal of this research effort is to apply deep learning to automatically detect and flag misclassified biometric images in large-scale datasets for subsequent examination and manual correction. Our primary focus is on intra-modality classification of fingerprints, specifically addressing 20 classes defined by finger identity (thumb, index, etc.) and collection method (flat vs. rolled). We utilize a modified Residual Network (ResNet) architecture in a Siamese configuration to generate robust image embeddings for comparison. Therefore, the specific objectives of this study are to (i) preprocess and organize fingerprint data into 20 distinct classes to create a viable dataset for testing; (ii) design and implement a Siamese network architecture fine-tuned for high-precision fingerprint classification; (iii) systematically test the network’s performance using a 3×4 full factorial design, analyzing the impact of different embedding dimensions (64, 128, 256, 512) and similarity thresholds (0.5, 0.2, 0.1) on classification metrics; and (iv) provide a proof-of-concept tool that quantifies the trade-off between increased conditional classification accuracy and the corresponding rise in the "uncertain" image classification rate, thus guiding practical application for error correction in operational datasets.

### Preliminary Work

The foundational concept for the automated biometric data classification tool originated from a preliminary system designed to handle multiple modalities, encompassing face, iris, and fingerprint images. A dedicated preprocessing module was developed to standardize inputs, converting raw WSQ files obtained from data collections into a 180 × 180 × 3 format compatible with the chosen neural network architecture. This initial system utilized a Residual Network (ResNet), pre-trained on the external ImageNet dataset, whose weights were then fine-tuned for biometric sorting over five epochs via transfer learning. Data used for both inter-modality (modality type) and intra-modality (within-modality errors) activities were sourced from numerous collections at the West Virginia University Biometrics Lab under approved IRB protocols. The specific intra-modality tasks included: classifying face images by pose (frontal, profile, other), iris images by side (left or right), and fingerprint images into 20 classes defined by both finger type (index, middle, ring, little, thumb) and collection methodology (flat or rolled impressions). A compilation of the initial results is summarized in Figure 1.

**Figure 1.**
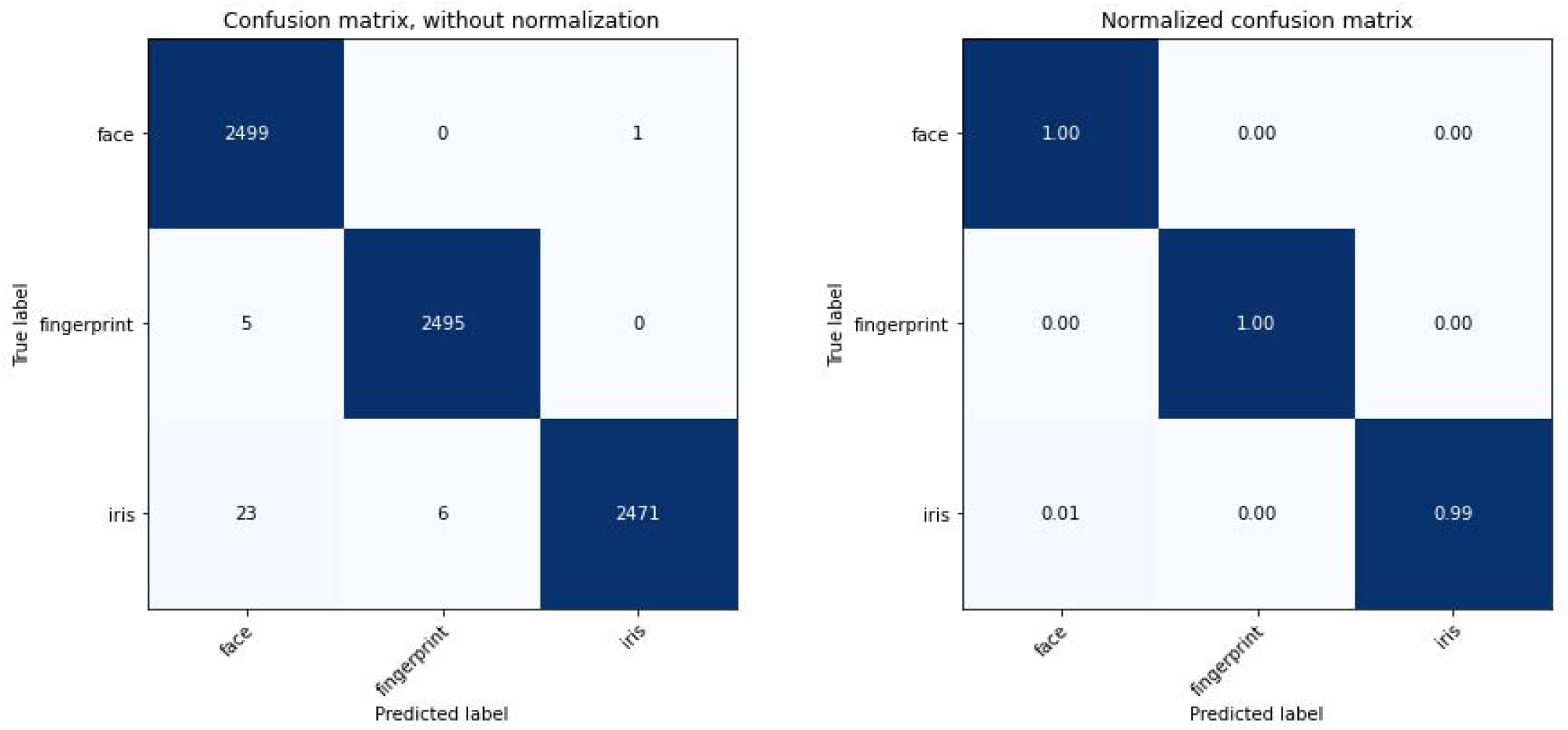
Face, Fingerprint and Iris Classification.

The system achieved exceptional performance on high-level tasks: the inter-modality classification of face/fingerprint/iris obtained an accuracy of 99%. Similarly, simple intra-modality sorting tasks, such as distinguishing left/right iris images and face pose classification, also yielded an accuracy of 99% (Figure 2). However, the crucial, fine-grained 20-class fingerprint classification task resulted in a markedly lower accuracy of 84. This performance deficit was critically analyzed and attributed to the network’s poor ability to extract sufficiently discriminative features to resolve minute differences between visually similar classes, causing significant confusion between sets like the left/right index and right/left thumbs. As illustrated by the Misclassified Fingerprints from Initial Testing (Figure 3), this high rate of misclassification for similar prints indicated the need for a network architecture specialized in metric learning. Consequently, the research pivot was made to a different neural network paradigm to improve the efficacy of fingerprint detection, specifically aimed at realizing the concept of a classification tool (Figure 4) that can automatically flag errors for necessary manual correction.

**Figure 2.**
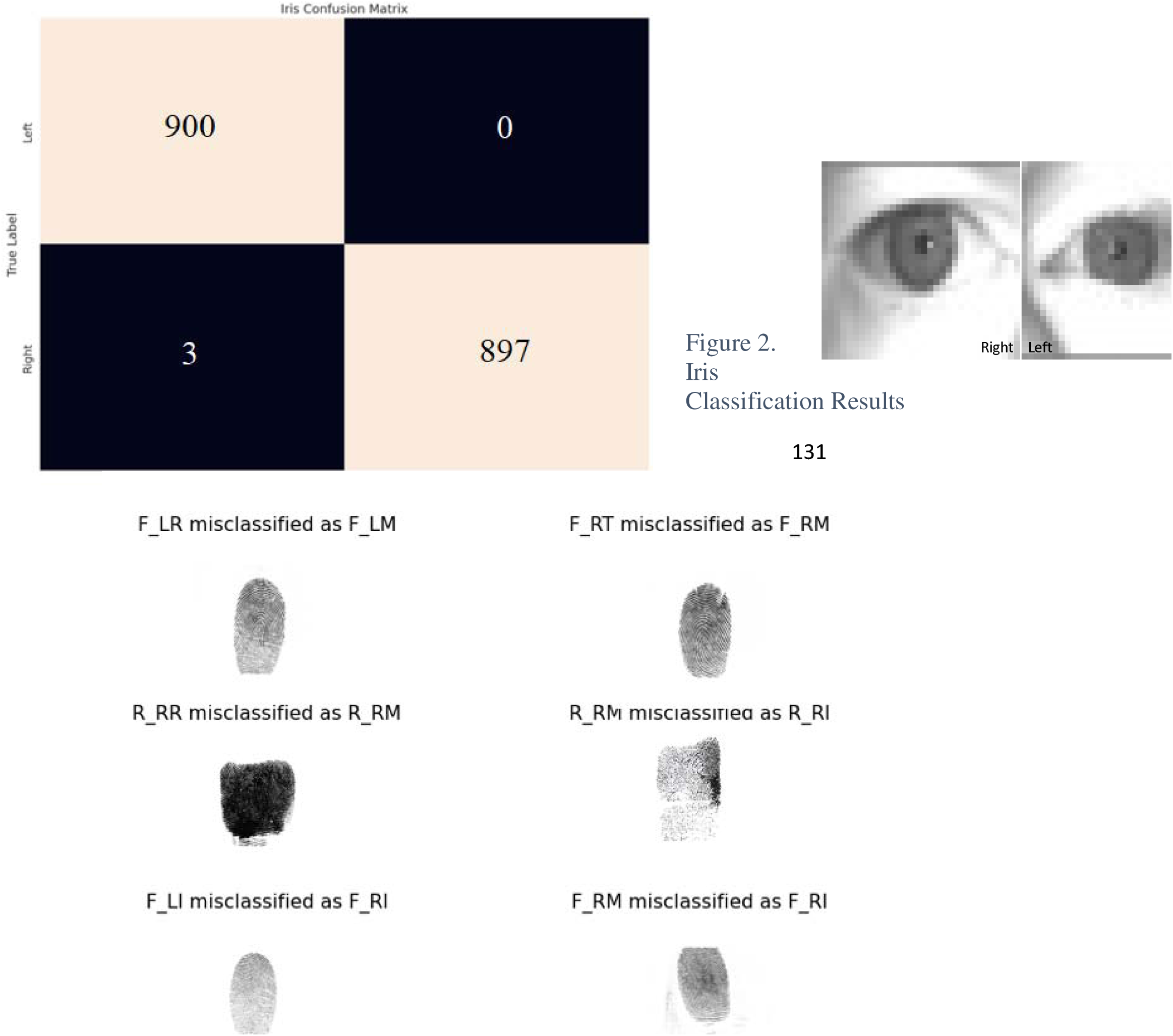
Iris Classification Results.

**Figure 3.**
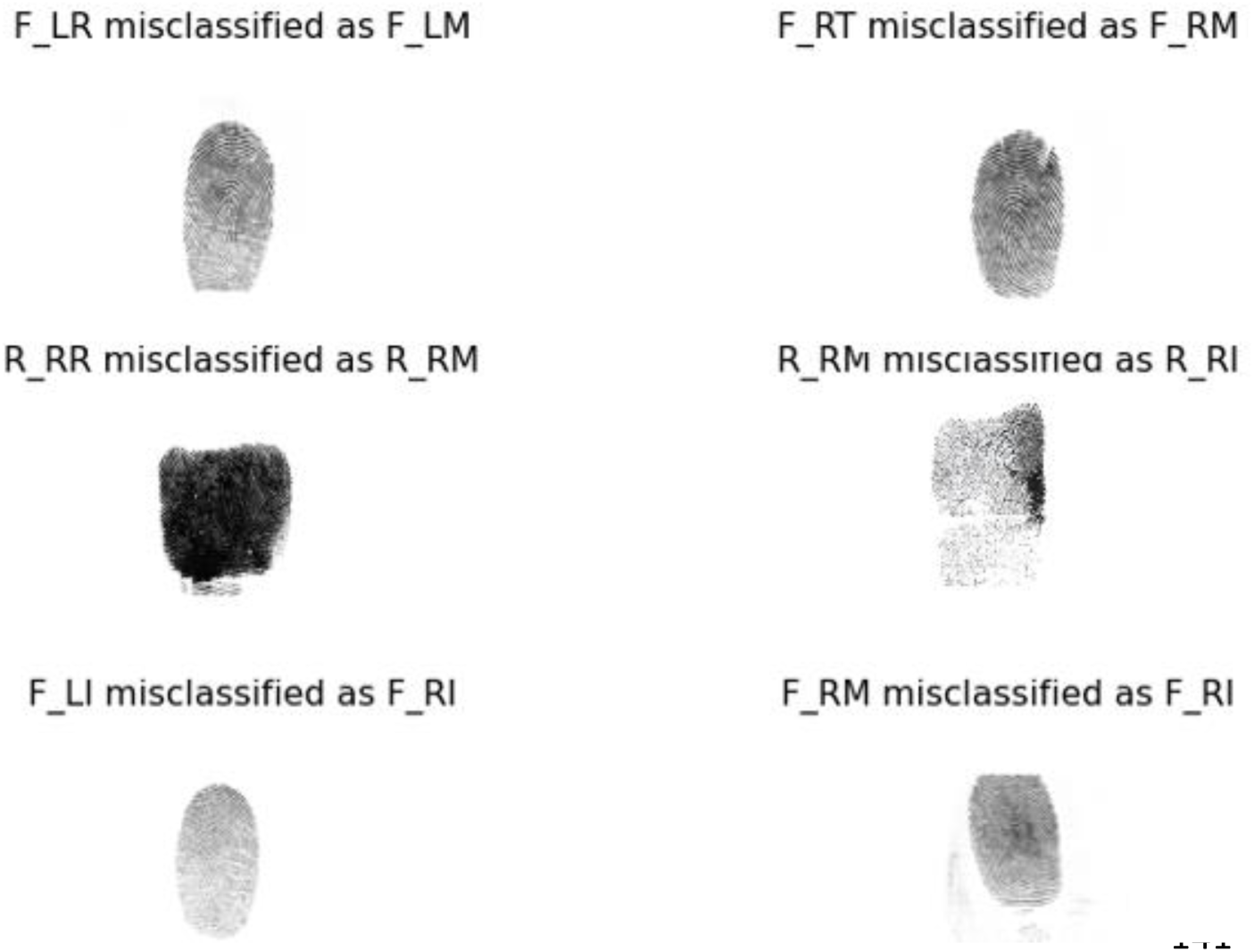
Misclassified Fingerprints from Initial Testing.

**Figure 4.**
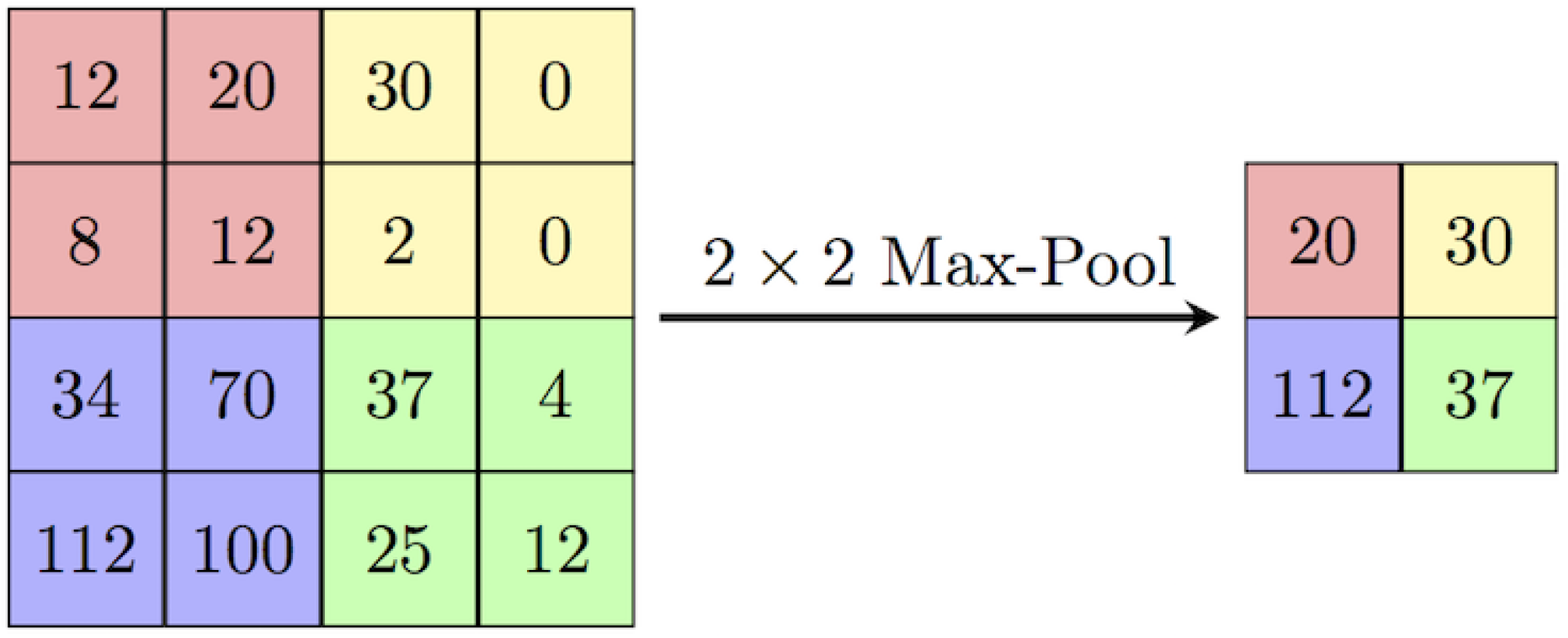
Max Pooling Demonstration [31].

## Methods

### A. Neural Networks Background

The field of Machine Learning (ML) constitutes an intersection of computer science, artificial intelligence, and statistics, fundamentally focused on developing computational systems that automatically improve performance through empirical experience [11]. Neural Networks (NNs) represent a specific, highly effective class of ML models, drawing conceptual inspiration from biological neural systems. These networks operate through interconnected nodes (neurons) that process and transform input data via an adjustable set of weights and biases, guided by algorithms. This concept, rooted in the principle of Hebbian learning that repeated activation strengthens neural connections [12] was first formalized computationally with the Perceptron in the 1950s. Although the field experienced an early recession, innovations in the 1980s, particularly the introduction of backpropagation and gradient descent, coupled with exponential advances in computing power (aligned with Moore’s Law), propelled NNs into a dominant paradigm [13]. In the contemporary era, NNs are essential for complex pattern recognition tasks across all sectors, making them highly pertinent for the specialized image classification required in biometric data analysis.

### B. Fingerprint Classification Background

Fingerprints represent one of the most reliable modalities in biometric systems, characterized by a unique sequence of ridges and furrows on the finger surface. The core structure is defined by distinct patterns such as the arch (ridges entering one side and exiting the other), loop (ridges curving and exiting the same side they entered), and whorl (ridges forming circular shapes) [14]. These macroscopic patterns are further detailed by minutiae, which are the local irregularities of the ridges, such as the ridge ending (where a ridge terminates) and the ridge bifurcation (where a ridge splits into two) [15]. The uniqueness and consistency of these patterns form the basis for identification, and fingerprint classification based on these feature types is a mature and well-understood field [16], with robust feature extractors developed to accurately capture salient characteristics from images [10].

Crucially, this research leverages the concept of human-level expertise in fingerprint analysis. Studies have demonstrated that expert latent print examiners, through sufficient training and experience, develop the ability to accurately distinguish between finger types (e.g., thumb vs. index) [17]. This trained human capability to sort prints based on their unique characteristics beyond core pattern types provides the direct conceptual foundation for our automated approach. By training a neural network to identify and correctly classify prints based on these distinct finger characteristics, we aim to replicate this expert sorting ability to identify subtle classification errors within large datasets [18].

### C. Image Classification

Image classification is a critical application area for modern machine learning, deeply relevant to this work’s objective of classifying biometric images. The standard architecture for analyzing and deciphering visual data is the Convolutional Neural Network (CNN) [19]. Unlike traditional multi-layer feed-forward networks, which struggle with the computational complexity of high-resolution image inputs, a CNN effectively manages large pixel densities by leveraging its architectural components to learn hierarchical features [20]. This is achieved by assigning learnable parameters, weights and biases, to automatically extract characteristics that distinguish one image class from another. The core of the CNN is the convolutional layer, which generates a feature map representing specific features extracted across all locations of the input image [19]. This localization and weight-sharing mechanism significantly reduces the number of free parameters compared to fully connected layers, enabling the network to scale to massive datasets. Following the convolutional layer, pooling layers are used to perform dimensionality reduction on the feature maps, optimizing computational efficiency and promoting robustness to minor spatial variations [21-22]. Two common methods are Max Pooling (which retains the maximum value within a defined kernel) and Average Pooling (which computes the average value within the kernel). This hierarchical feature extraction makes CNNs highly effective and computationally feasible for high-accuracy classification tasks, such as those required for fingerprint analysis (Figure 7).

**Figure 5.**
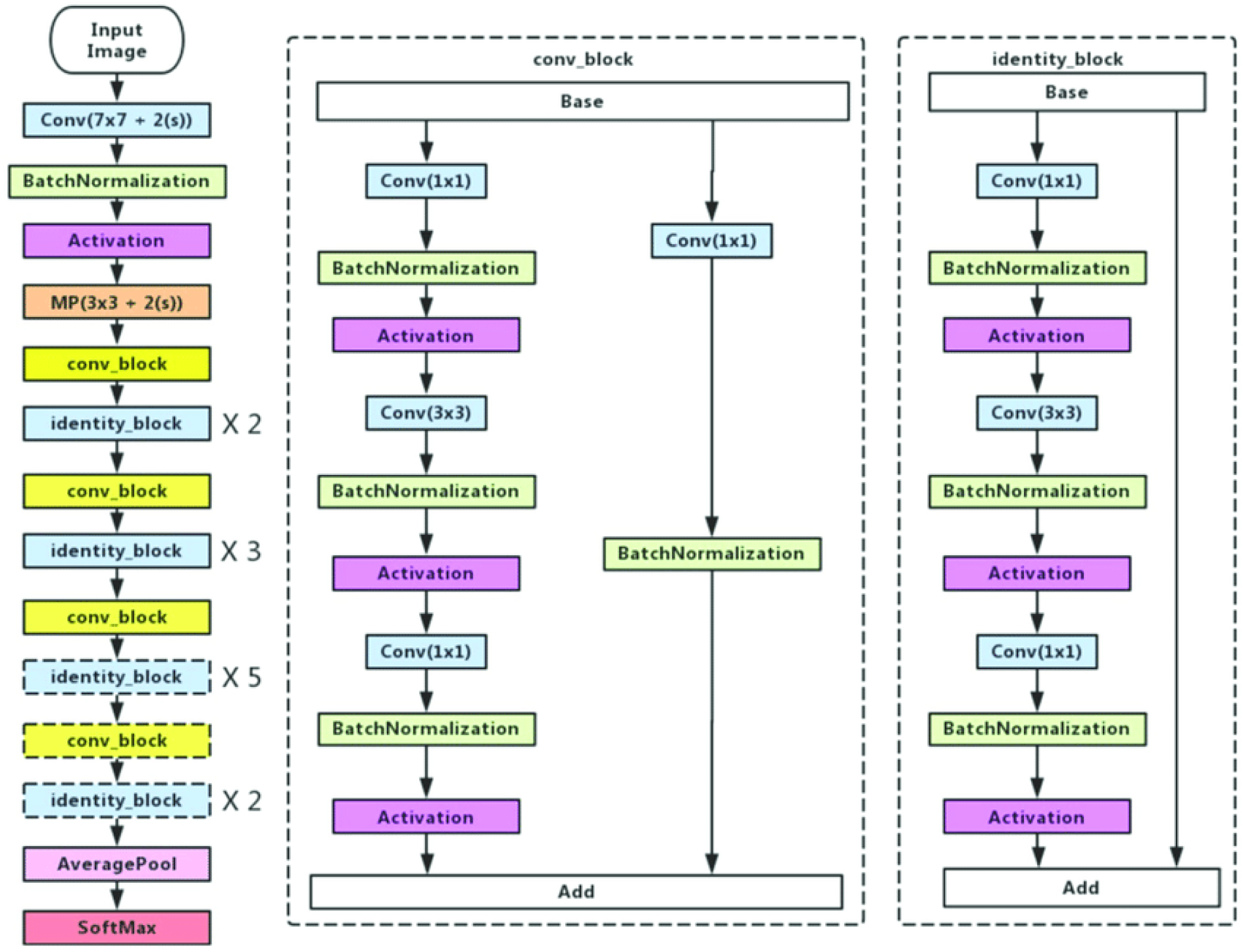
Resnet50 Architecture [24].

**Figure 6.**
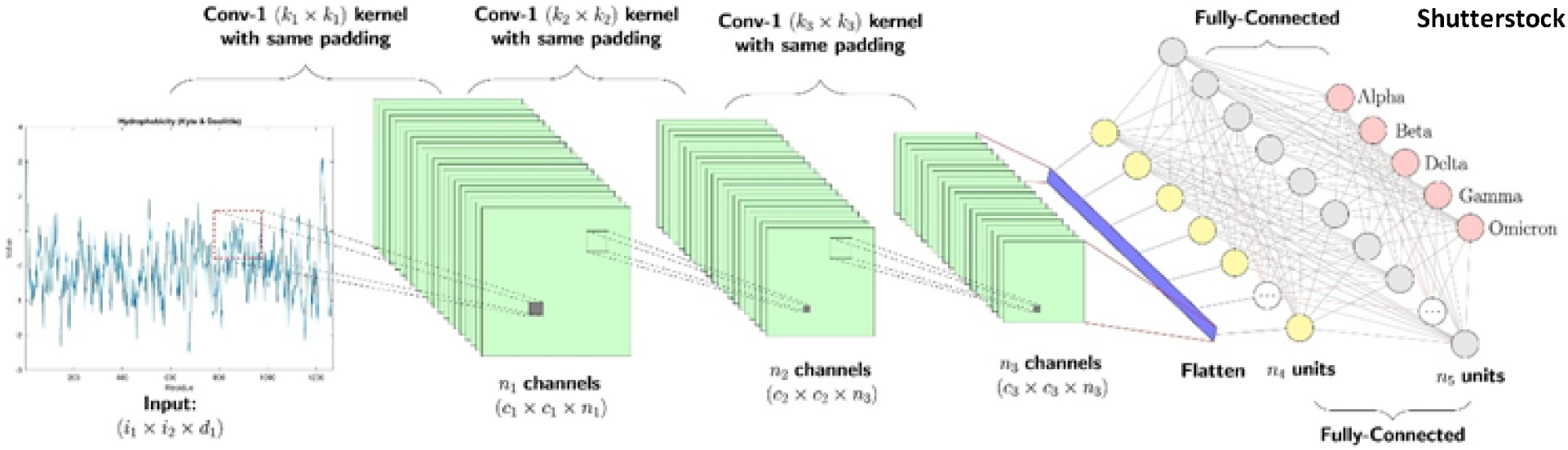
Siamese network architecture.

**Figure 7:**
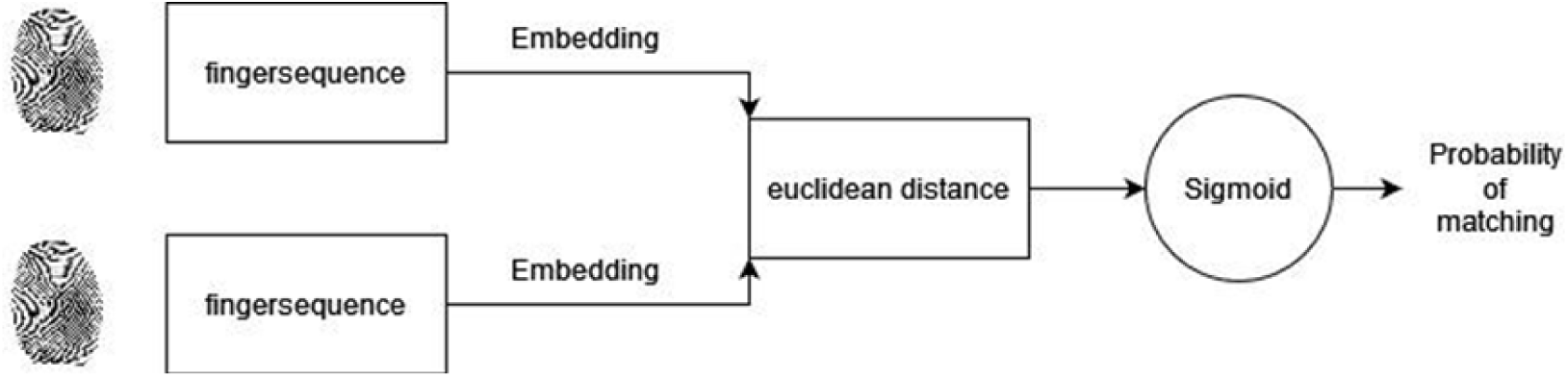
Siamese Network Architecture.

With the rapid progress in deep learning, advanced architectures have propelled image classification to new levels of performance and applicability [23]. For the purposes of this research, we focus on two of the most recognized and influential Convolutional Neural Network (CNN) architectures: ResNet and the Siamese Network. The Residual Network (ResNet) family, particularly ResNet50, is celebrated for its ability to train models with hundreds of layers while maintaining high accuracy, primarily through the innovative use of skip connections. This make ResNet a robust base for complex feature extraction in large biometric datasets. This architecture provides the necessary foundation for the subsequent implementation of the Siamese network, which is specifically designed for metric learning essential to our classification and error detection task.

### D. ResNet50 Architecture

ResNet50 is a specific Residual Network architecture introduced by Microsoft Research that achieved distinction by winning the ImageNet Large Scale Visual Recognition Challenge (ILSVRC) for its classification efficiency on massive datasets (Figure 8) [21]. The primary innovation of ResNet is its ability to successfully train models with depths ranging from hundreds to thousands of layers while maintaining, and even increasing, accuracy. Historically, merely adding more layers to a standard Deep Convolutional Neural Network (CNN) led to the problem of degradation, where accuracy would saturate and then rapidly decrease due to convergence difficulties and optimization problems [24].

**Figure 8.**
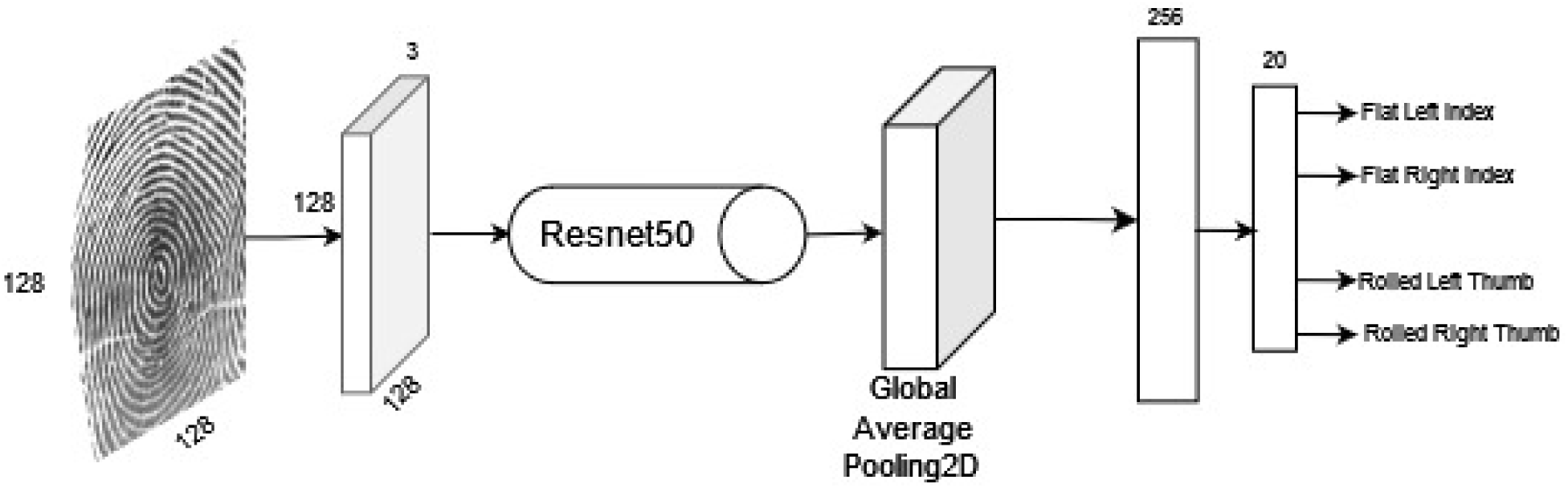
An initial architecture.

Residual Networks solve this critical challenge through the use of skip connections (or identity mappings). Instead of forcing the network to learn the entire function layer by layer, a skip connection creates an alternate shortcut that bypasses one or more layers, allowing the output from a previous layer to be added directly to the output of a later stacked layer. This formulation ensures that the network is learning the *residual mapping* rather than the original complex mapping. This "skipping" mechanism facilitates easier gradient flow during backpropagation and enables deeper layers to perform at least as well as their shallower counterparts. This architectural efficiency makes ResNet an ideal feature extraction backbone for applications like our fingerprint classification, where deep learning is used to efficiently sort through massive datasets and label misclassifications.

### E. Siamese Network Architecture

The Siamese network, a class of neural architectures introduced in the 1990s [25], is specifically designed for similarity learning (or metric learning). It consists of two or more identical subnetworks that share the same configuration, parameters, and weights.

This structure allows both subnetworks to generate feature vectors (or embeddings) for their respective inputs, which are then compared. Siamese networks are commonly applied to problems such as verification, for instance, determining if two input face images belong to the same individual (Figure 9). In the context of this paper, the same verification principle is applied to the classification of fingerprints [26]. The network is trained using input pairs: a designated anchor image is compared against either a genuine pair (an image belonging to the same finger class) or an impostor pair (an image belonging to a different finger class). The anchor image essentially functions as a representative for its specific class. By comparing the anchor’s embedding against all other prints in a dataset, the network generates a quantifiable distance metric that estimates the closeness or distance of other prints to that representative class, enabling fine-grained class separation. This comparative training mechanism is used to determine whether a given print matches the anchor’s identity.

**Figure 9.**
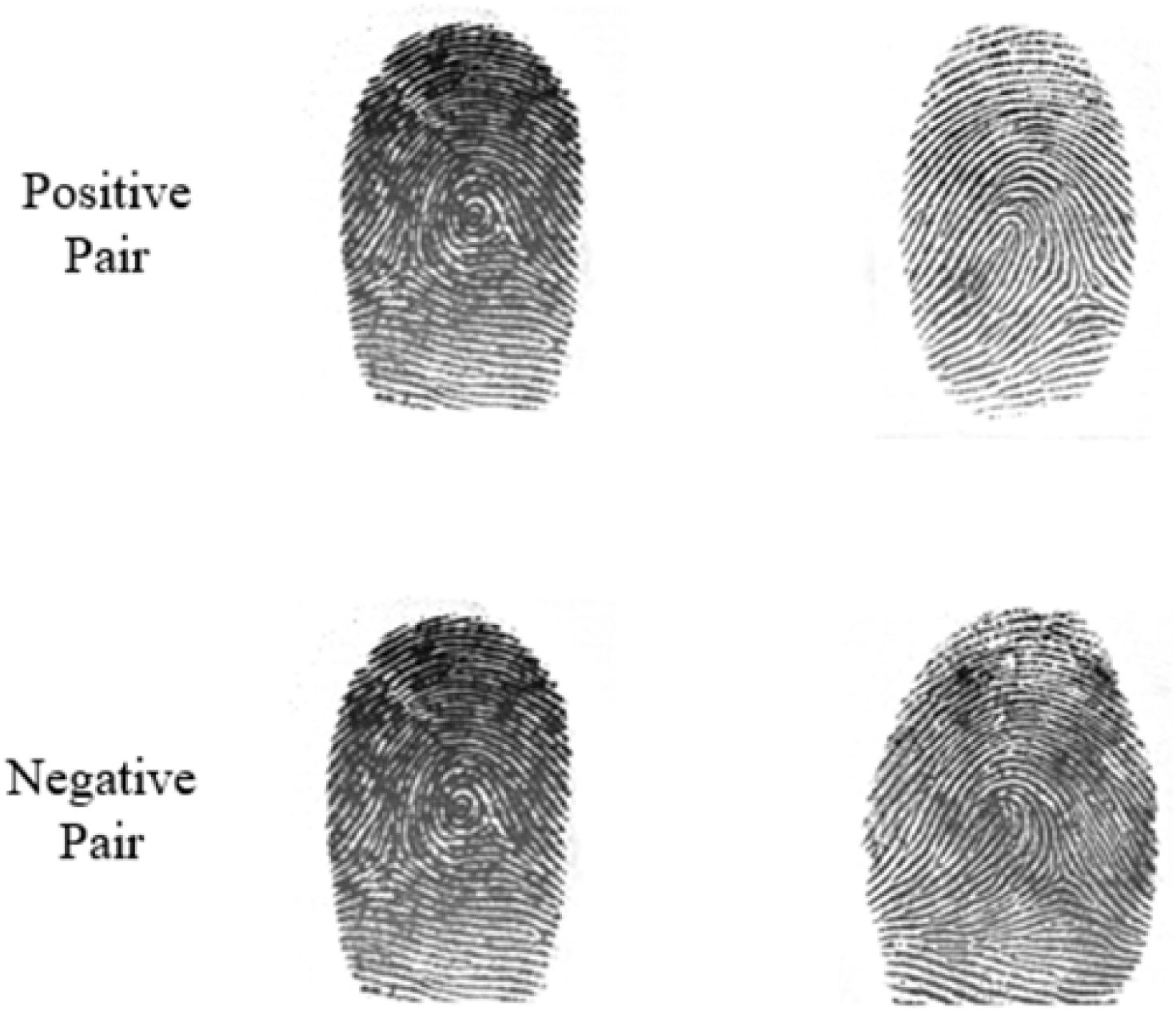
Example of Positive Pair (Index and Index) with Negative Pair (Index and Thumb)

## Metrics

### F. Outcome Metrics

Outcome metrics are a means to quantitatively assess the results of the experiment. In this work, various outcome metrics were used to assess the usability of the neural network.

#### i) Accuracy

Accuracy is a metric that allows to evaluate our effectiveness for classification models. It is the number of correct predictions over the total number of predictions [27].

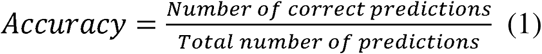

Accuracy is calculated by positive and negative terms. There is True Positive (TP) where the model predicts the positive class correctly. True Negative (TN) is where the model predicts the negative class correctly. False Positive (FP) is when the model predicts a class to be correct when it is not. False Negative (FN) is when the model predicts a class to be incorrect when it is not. By combining these terms, it is possible to obtain a numerical value for accuracy:

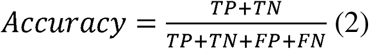

In the case of this research, accuracy defines the number of fingerprints correctly classified over the total number of fingerprints in the dataset.

#### 2) Conditional and Unconditional Accuracy

In the case of this research, there are different ways of calculating the accuracy as it is not simple as the case of only positives and negatives. In order to provide a full picture, we calculated conditional and unconditional accuracy. Conditional accuracy only considers the images that are classified as positive or negative but does not include the images that are uncertain. Unconditional accuracy on the other hand considers all images/factors.

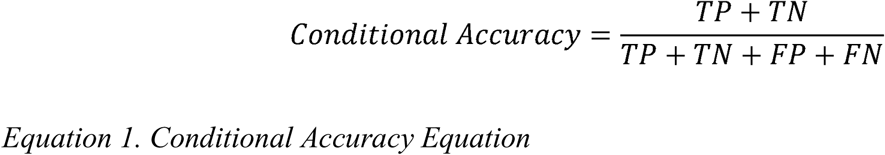

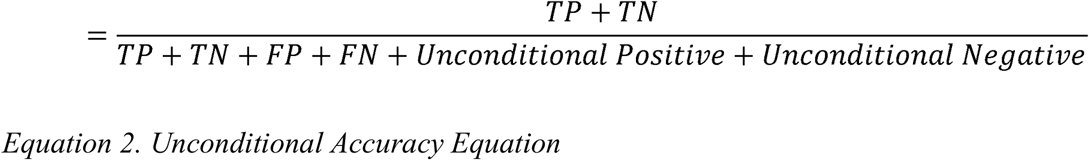

#### 3) Uncertain

Uncertain are predictions that fall in the range of threshold and 1-threshold. These are predictions for which there is not enough sufficient confidence to mark them as positive or negative. These can be images that have been tampered with (blurred or lower quality) or have some sort of error that the neural network is unable to classify accurately.

#### 4) Precision

Precision determines which proportion of positive identifications were accurately predicted to be correct. It is when True Positive is over the combination of True Positive and False Positive [28].

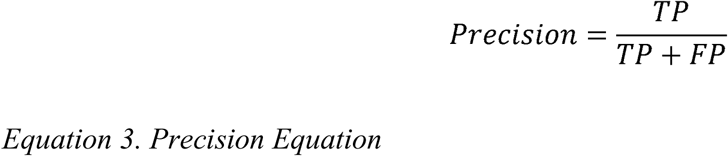

In the case of this research, precision defines the network’s ability to correctly predict and classify specific classes of the fingerprints, such as front left index and other classes.

#### 5) Recall

Recall determines which proportion of actual positive identifications were correct. It is calculated using True Positive over the combination of True Positive and False Negative [28].

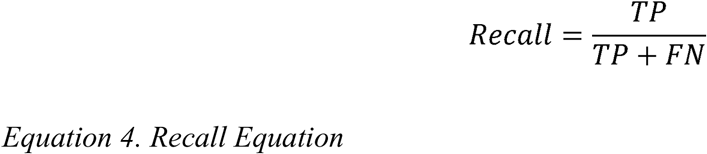

#### 6) Conditional and Unconditional Recall

In the case of this research, there are different ways of calculating the recall as it is not simple as the case of only positives and negatives. In order to provide a full picture, we calculated conditional and unconditional recall. Conditional recall only considers the images that are classified as positive or negative but does not include the images that are uncertain. Unconditional recall on the other hand considers all images.

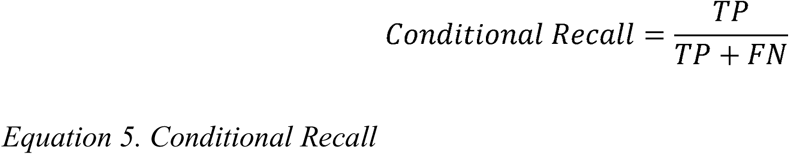

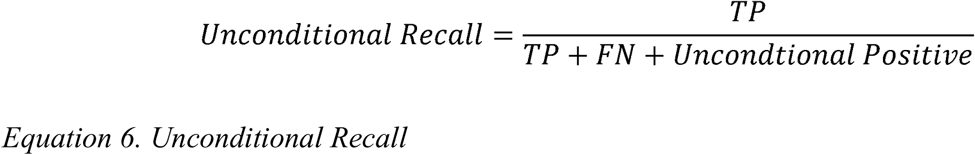

#### 7) F1 Score

The F1 Score, also known as the F-measure, is a metric which is based on error. It measures the neural network model’s performance by calculating the harmonic mean of precision and recall for the minority positive class [29]. It is one of the most commonly used metrics for classification models as it provides easy to understand results for balanced and imbalanced datasets factoring in the precision and recall values.

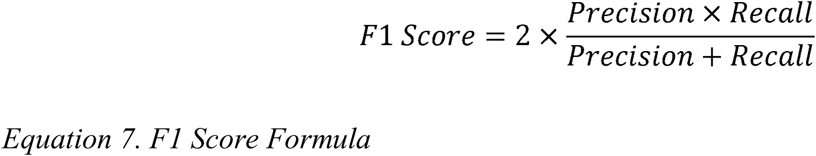

To interpret the score, F1 provides an overall model performance from 0 to 1, with 1 being the best possible score. It shows the model’s ability to detect positive cases in recall and accurately classified cases in precision. In the scope of this research, there will be Conditional F1 and Unconditional F1, with one considering the uncertain factor while the other does not (Table 1).

**Table 1.** F1 Score Distribution.

| F1 Score | Interpretation of Score |
| --- | --- |
| Greater than 0.9 | Excellent |
| 0.8 – 0.9 | Great |
| 0.5 – 0.8 | Average/Medium |
| Less than 0.5 | Low/Undesirable |

### Experiment

#### Initial Baseline Classification using ResNet50

For the biometric data classification effort described here, all raw fingerprint files in the WSQ format were preprocessed and uniformly converted into an 8-bit, 128 × 128 × 3 PNG image structure. The initial classification was centered on a Residual Network (ResNet50) architecture. Network weights were initialized via transfer learning from a model pre-trained on the ImageNet Large Scale Visual Recognition Challenge (ILSVRC) dataset, which encompasses the classification of 1,000 diverse object categories [3]. The transfer learning process involved fine-tuning these weights for the specific biometric sorting purposes, utilizing 5 training epochs.

The data utilized was divided into two distinct sets (i) Inter-modality classification was trained and tested on a dedicated, small subset of operational data, comprising 7,500 images (2,500 for each modality: face, iris, and fingerprint). (ii) Intra-modality activities (the primary focus) used a large proprietary dataset collected by the West Virginia University Biometrics Lab under approved IRB protocols, consisting of 40,353 images for training and 10,088 images for testing. The fingerprint images for the intra-modality task were uniformly processed to grayscale and organized into 20 distinct classes. These classes are defined by the cross-product of the 10 unique finger identities (left/right thumb, index, middle, ring, little) and the two collection methodologies (flat or rolled impressions).

#### ResNet50 Architecture and Baseline Performance

The initial 128 × 128 × 3 image input was passed through the ResNet50 network. The network’s core feature extraction was accomplished by the Global Average Pooling layer, which computes the average feature map value across the spatial dimensions. This extraction pipeline then fed into a dense layer of 256 neurons, which served as the image embedding (a reduced-dimensionality vector representation). The final layer was a classification head consisting of 20 neurons, each corresponding to one of the 20 possible finger classes. The network’s output used a one-hot encoding scheme, setting only one output neuron to 1 and the rest to 0 to indicate the predicted class (e.g., a "flat left index" image would correspond to its specific class neuron). The overall classification accuracy for this initial ResNet50 network was 87%. While adequate for simple image classification, this accuracy level was deemed insufficient for reliably scouring large operational datasets where a high degree of certainty is required for flagging misclassified records, thus necessitating an improved network architecture.

To achieve more robust and accurate classification performance, the base ResNet50 feature extractor was adapted into a Siamese network configuration, as conceptually illustrated in Figure 10.

**Figure 10:**
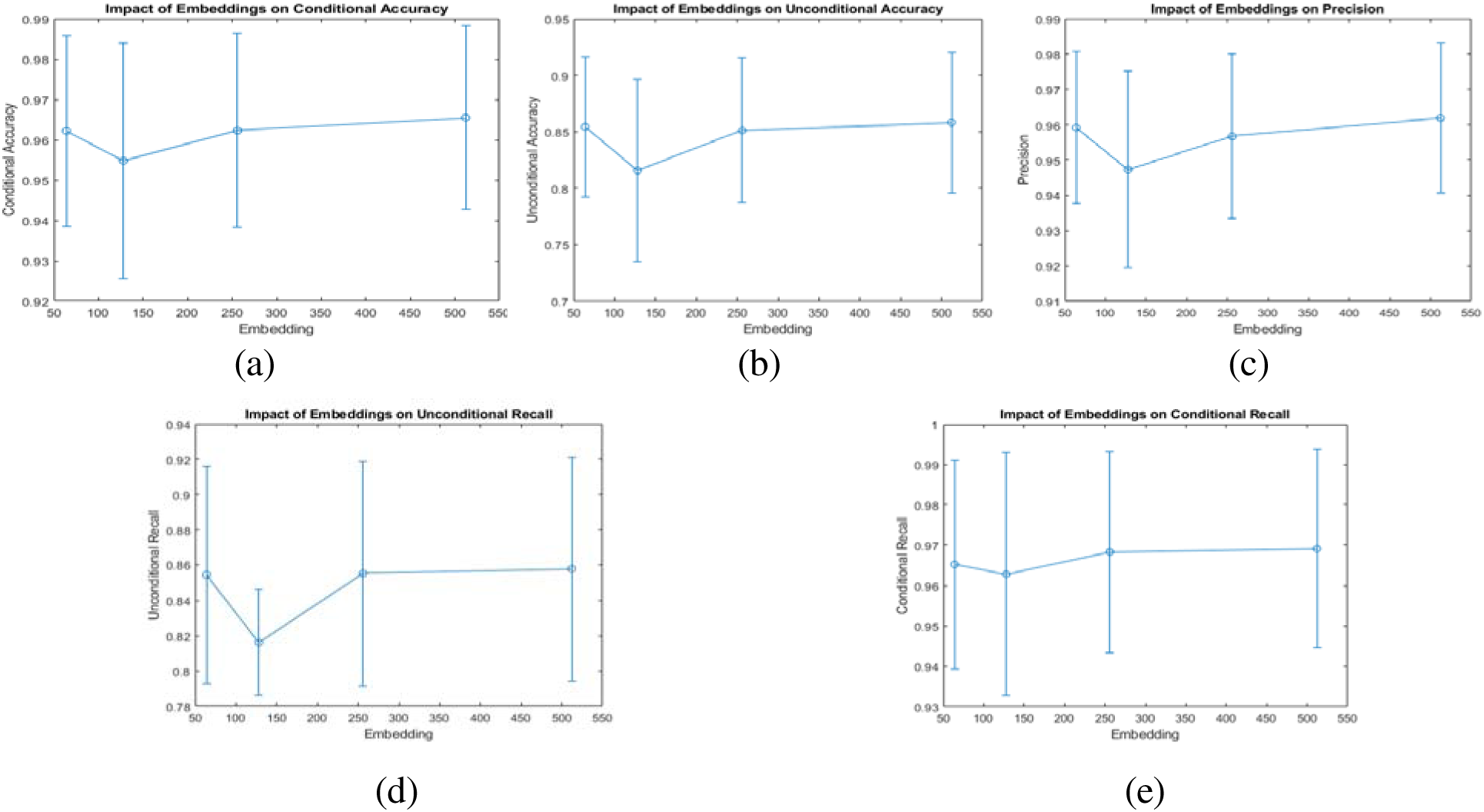
Impact of Embeddings on Different Metrics: (a) Conditional Accuracy, (b) Unconditional Accuracy, (c) Precision, (d) Unconditional Recall, and (e) Conditional Recall.

This architecture consists of two identical subnetworks, each acting as a weight-sharing "finger sequence network." Each subnetwork is modified to produce a 256-dimensional embedding vector for its respective input image. The primary function of these embeddings is to serve as low-dimensional vector representations that encode the salient features of the image, thus reducing data dimensionality while preserving discriminative information. Following the feature extraction, the two embedding vectors are passed to the Euclidean Distance layer. This metric serves as a distance measure, quantifying the relative dissimilarity between the two inputs by calculating the L2 norm between the two 256-dimensional vectors. This distance value is then fed into a sigmoid function. The sigmoid output produces a probability of matching ranging between 0 and 1, where a value near 0 indicates high similarity (the images belong to the same class) and a value near 1 indicates low similarity (the images belong to different classes or classes).

The initial network architecture relied on a 20-dimensional output vector where each element was mapped to a specific classification class. In contrast, the Siamese network adopts a metric learning paradigm and is trained using input pairs, as depicted in Figure 11. A representative image, referred to as the anchor, is fed into one side of the network. The network’s training objective is to learn an embedding space where the genuine pair (an image of the same class as the anchor) yields a distance close to 0, and the impostor pair (an image that is not of the same class as the anchor) yields a distance close to 1. Specifically, the network is trained with positive pairs, where the second image belongs to the same class as the anchor, and negative pairs, where the second image belongs to a different class (Figure 12). This comparative training mechanism forces the network to create highly discriminative embeddings that maximize the separation between inter-class feature vectors while minimizing the distance between intra-class feature vectors.

**Figure 11:**
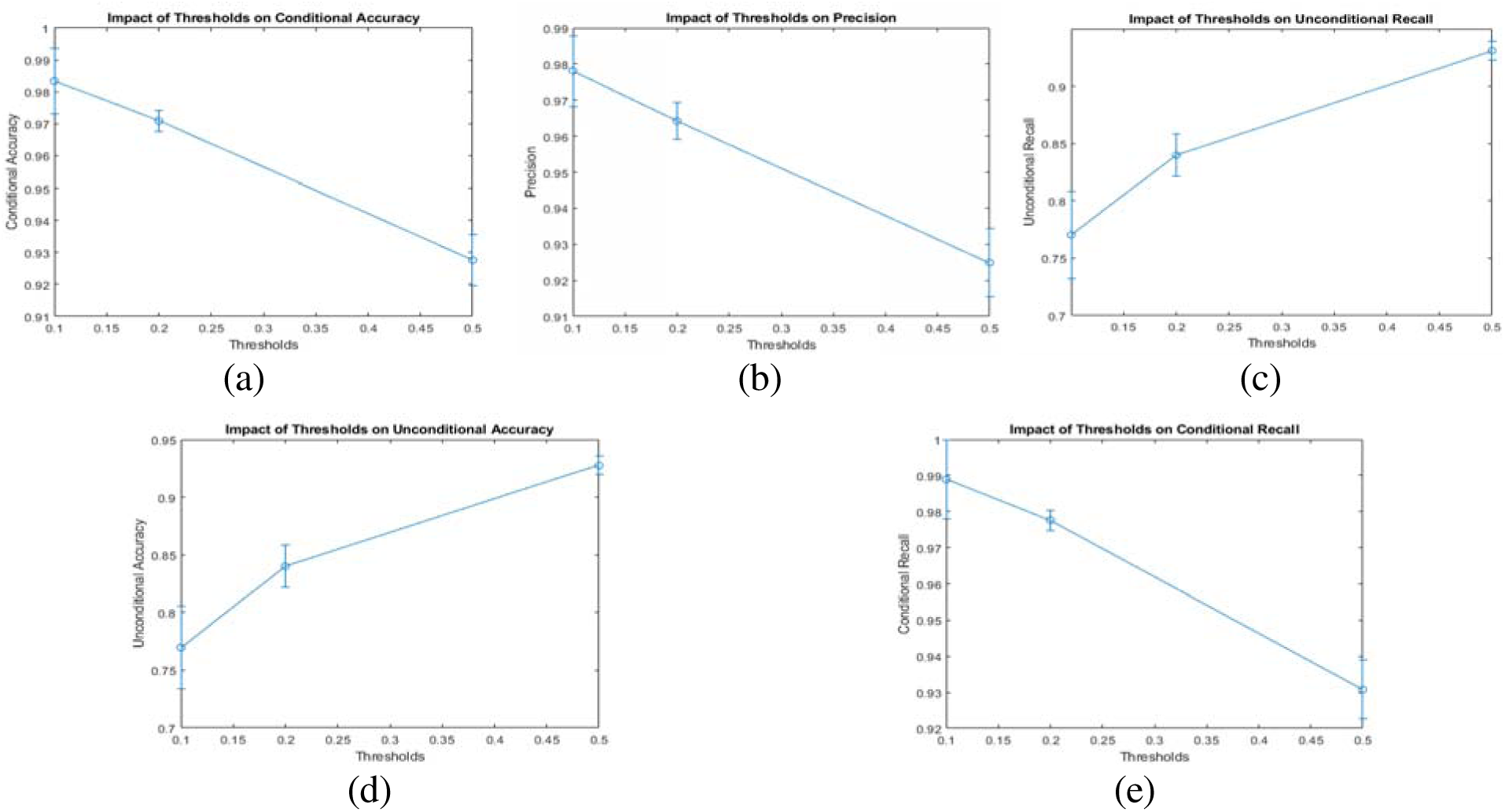
Impact of Thresholds on Different Metrics: (a) Conditional Accuracy, (b) Precision, (c) Unconditional Recall, (d) Unconditional Accuracy, (e) Conditional Recall.

**Figure 12.**
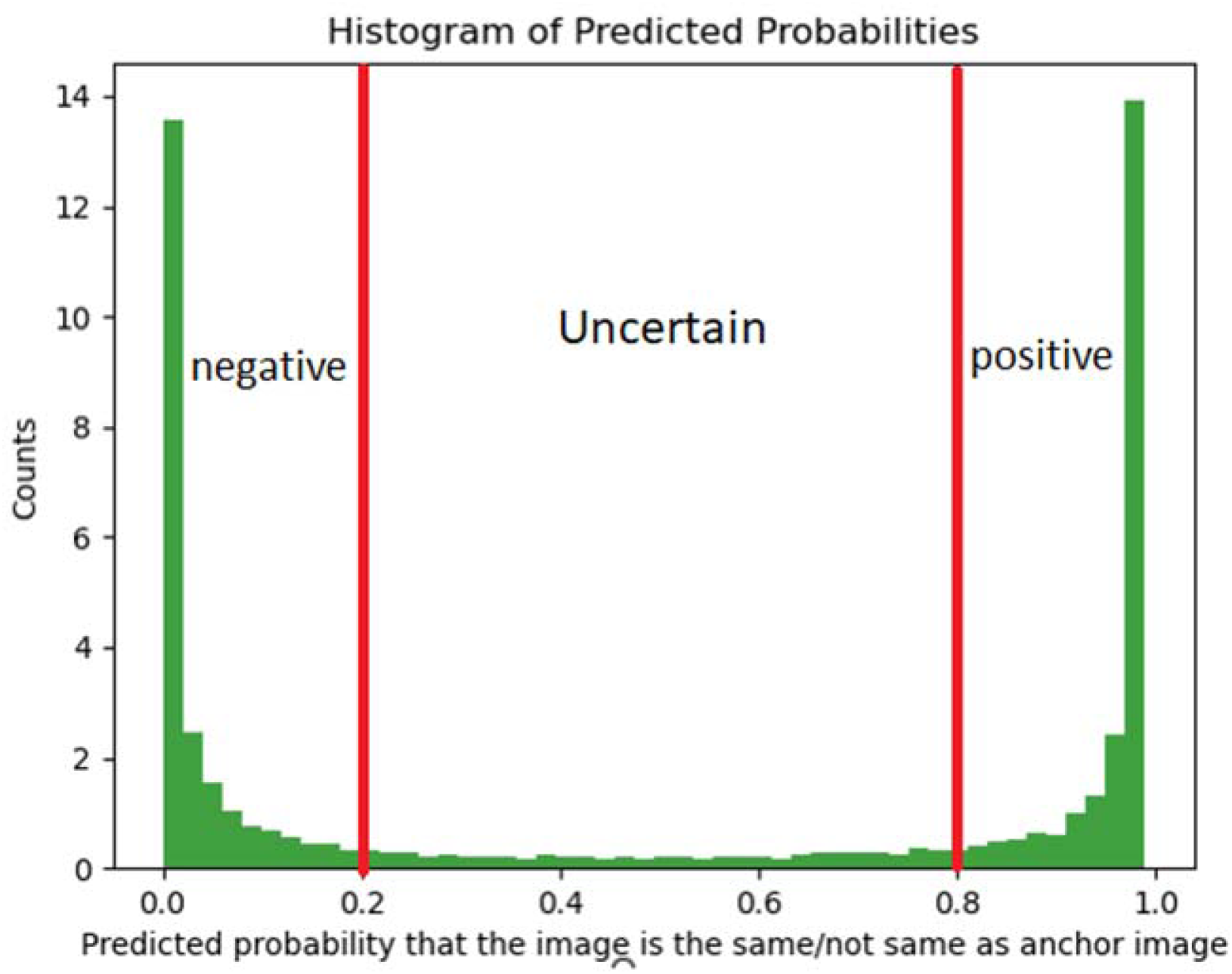
Combined Histogram of Predicted Probabilities of Results.

In this study, a 3 × 4 full factorial experimental design was implemented to rigorously evaluate the sensitivity of the Siamese network’s performance to two key hyper-parameters: the embedding dimension and the similarity threshold. This systematic approach allowed us to quantify the impact of these components on the network’s precision, conditional accuracy, unconditional accuracy, conditional recall, unconditional recall, and the resulting uncertainty factor. Specifically, we tested four embedding dimensions (64, 128, 256, and 512) against three similarity thresholds (0.1, 0.2, and 0.5). The inclusion of both the conditional and unconditional variants of accuracy and recall was crucial to precisely determine the magnitude of the "uncertain factor", the percentage of images that fall within the threshold boundary and require manual intervention. To ensure statistical rigor and mitigate the effect of random initialization, each experimental condition was replicated 10 times. The resulting mean values and standard deviations are comprehensively summarized in Figure 13 and Figure 14.

The analysis of the experimental results demonstrates the differential impact of the tested hyper-parameters. Varying the embedding dimensions (64, 128, 256, and 512) had no statistically significant impact on classification accuracy or the other evaluated metrics, as evidenced across Figure 2 (a-e). This suggests that a 64-dimensional embedding is sufficient for feature discrimination within this specific metric space. In sharp contrast, modifying the similarity threshold (using subsets of 0.1, 0.2, and 0.5) yielded a significant and systematic change in performance. Narrowing the threshold (i.e., decreasing the value from 0.5 to 0.1) consistently resulted in a substantial increase in conditional accuracy and recall (Figure 14 (a-e)). Crucially, this improvement simultaneously increased the uncertain factor from 0% (at a 0.5 threshold) to 24% (at a 0.1 threshold). The uncertain factor explicitly represents the number of image comparisons where the Euclidean distance falls within the range of [Threshold, 1 - Threshold]. These are predictions for which the network lacks sufficient confidence to assign a definitive positive or negative classification, consequently requiring manual review. The impact of this factor is clearly visible when comparing the conditional versus unconditional metrics of accuracy and recall (Figure 13 (a) and (e)), where the introduction of the uncertain images drastically depresses the unconditional values. This systematic dependency means that while smaller threshold values yield higher certainty in the classifications made, they dramatically increase the volume of images labeled as "uncertain" (Figure 15). This creates a direct operational trade-off: achieving a high conditional classification accuracy (e.g., 98%) demands accepting an elevated uncertain rate (e.g., 24%) (Table 2). Enhancement of fingerprint image using multiple filters has also been demonstrated [30]. For practical application in databases containing millions of records, a 24% uncertainty level translates to an immense number of prints that must be manually inspected or passed through more sophisticated, yet time-consuming, secondary classification algorithms. Therefore, successful deployment of this proof-of-concept network requires careful optimization to balance the requirement for high classification rigor against the operational feasibility of managing the resulting uncertainty volume. The combination of a high-quality benchmark database and an innovative assessment approach are considered to be a strong foundation for future research, contributing to the development of more reliable and secure biometric systems [31]. Fingerprint matching for noisy and distorted patterns using a Siamese Network with ResNet50 and multihead attention has been reported [32]. FootprintNet, a Siamese network that utilizes pre-trained convolutional neural networks, specifically EfficientNet, MobileNet, and ShuffleNet, to improve the robustness and accuracy of footprint recognition has recently been proposed [33]

**Table 2.** Summary of results of threshold, accuracy, uncertain and recall relationship.

| Threshold | Accuracy | Uncertain | Recall/True Positive Rate |
| --- | --- | --- | --- |
| 0.5 | 92.5% | 0% | 92.9% |
| 0.2 | 96.8% | 14% | 96.4% |
| 0.1 | 98.4% | 24% | 98.0% |

## CONCLUSION

Fingerprints are indispensable for identification in modern biometrics, yet the integrity of large-scale databases is continuously threatened by erroneous classification, a critical concern where a single mislabeled image can compromise an entire archive. To address this, we successfully developed a Siamese network for the precise classification of fingerprints by finger type and collection methodology (flat versus rolled impressions). A central objective was to analyze the practical utility of this tool by systematically testing the effect of various embedding dimensions and similarity thresholds (specifically 0.5, 0.2, and 0.1) on performance metrics. Our primary finding reveals a crucial trade-off for operational deployment: using a lower similarity threshold significantly boosts conditional classification accuracy (achieving up to 98%) and precision, but this comes at the expense of classifying a much larger percentage of images as "uncertain" (up to 24). This forced manual review of millions of uncertain records in large datasets highlights the challenge of balancing optimal statistical accuracy with practical operational efficiency and cost. This research delivers a robust proof-of-concept tool that moves beyond pure statistical accuracy to provide a quantifiable, practical method for identifying and flagging data integrity errors. The framework is modality-agnostic, making it suitable for deployment across face, iris, and other biometric data. Ultimately, this work contributes to establishing a standardized preprocessing procedure, a lightweight and efficient system that can be applied before any large biometric dataset is finalized, ensuring the accuracy and legitimacy of identity records while concurrently achieving substantial savings in time and operational costs.

## Author Contributions

JMD and TIS conceived idea; TIS, design, conducted experiments, interpreted data and prepared manuscript; JMD and NMN, supervised, reviewed and improved the manuscript; All authors agreed to submit the manuscript.

## Acknowledgements

TIS is thankful to the funding GRA provided by the JMD during this study.

## Data Availability Statement

All data are included in this manuscript.

